# Serotonin and neuropeptides are both released by the HSN command neuron to initiate *C. elegans* egg laying

**DOI:** 10.1101/323857

**Authors:** Jacob Brewer, Kevin M. Collins, Michael R. Koelle

**Author notes:** Corresponding author E-Mail (MRK).

## Abstract

Neurons typically release both a small molecule neurotransmitter and one or more neuropeptides, but how these two types of signal from the same neuron might act together remains largely obscure. For example, serotonergic neurons in mammalian brain express the neuropeptide Substance P, but it is unclear how serotonin signaling might be modulated by a coreleased neuropeptide. We studied this issue in *C. elegans*, in which all serotonergic neurons express the neuropeptide NLP-3. The serotonergic Hermaphrodite Specific Neurons (HSNs) are command motor neurons within the egg-laying circuit that have previously been shown to release serotonin to initiate egg-laying behavior. We found that egg-laying defects in animals lacking serotonin were far milder than in animals lacking HSNs, suggesting that HSNs must release other signal(s) in addition to serotonin to stimulate egg laying. While null mutants for *nlp-3* had only mild egg-laying defects, animals lacking both serotonin and NLP-3 had severe defects, like those of animals lacking HSNs. Optogenetic activation of HSNs induced egg laying in wild-type animals, or in mutant animals lacking either serotonin or NLP-3, but failed to induce egg laying in animals lacking both. We recorded calcium activity in the egg-laying muscles of animals lacking either serotonin, NLP-3, or both. The single mutants, and to a greater extent the double mutant, showed muscle activity that was uncoordinated and unable to expel eggs, such that the vm2 muscles cells that are direct postsynaptic targets of the HSN failed to contract simultaneously with other egg-laying muscle cells. Our results show that the HSN neurons use serotonin and the neuropeptide NLP-3 as partially redundant cotransmitters that together stimulate and coordinate activity of the target cells onto which they are released.

## Author Summary

Activity of the brain results from neurons communicating with each other using chemical signals. A typical neuron releases two kinds of chemical signals: a neurotransmitter, such as serotonin, and one or more small proteins, called neuropeptides. For example, many human brain neurons that release serotonin, a neurotransmitter thought to be involved in depression, also release the neuropeptide Substance P. Neuroscientists have typically studied the effects of neurotransmitters and neuropeptides separately, without considering how a neuron might use the two types of signals together. Here we analyzed how specific neurons in the model organism *C. elegans* use both serotonin and a neuropeptide together. The Hermaphrodite Specific Neurons (HSNs) activate a small group of neurons and muscles to generate egg-laying behavior. Killing the HSNs resulted in animals unable to lay eggs, but we found that eliminating either serotonin or the neuropeptide resulted in HSNs that still remained able to activate egg laying. However, eliminating both serotonin and the neuropeptide resulted in HSNs unable to activate coordinated contractions of the egg-laying muscles. Our results show that in a living animal, serotonin acts in concert with a coreleased neuropeptide to carry out its functions.

## Introduction

Drugs that selectively manipulate serotonin signaling are widely used to treat depression and other psychiatric disorders, yet these drugs are often ineffective, and no specific molecular defects in serotonin signaling have been identified as the cause of these disorders (1). This situation suggests there is more to understand about the basic science of serotonin signaling that could help explain the cause of many psychiatric disorders. One feature of serotonin signaling in the mammalian brain that remains poorly understood is that serotonin neurons appear to also release a specific neuropeptide, Substance P (2–6). Of the ~80 billion neurons in the human brain, only about 100,000 make serotonin: their cell bodies are concentrated in the raphe nuclei of the brain stem, but extend axons throughout the brain that release serotonin to influence many brain functions (2,7,8). Several methods have been used to measure the proportion of serotonin neurons that express Substance P in the various raphe subnuclei of human or rat brain, with results suggesting that from 25% to nearly all serotonin neurons also express substance P (2,5,9,10). The apparent co-release of serotonin and Substance P from the same neurons is just one instance of the broad but poorly studied phenomenon of cotransmission by small-molecule neurotransmitters and neuropeptides (11–14). It remains unclear how exactly coreleased serotonin and neuropeptide might functionally interact. Clinical studies of Substance P antagonists showed that they, like selective serotonin reuptake inhibitors, can have significant anti-depressant activity (15–17). One study in the brain stem respiratory circuit indicated that serotonin and substance P each independently stimulate activity of the circuit (10), but the complexity of mammalian brain circuits makes precise analysis of such effects difficult.

Small neural circuits of invertebrates provide the potential for more precise analysis of the functional effects of cotransmission. In such circuits, every cell can be identified, the small molecule neurotransmitter and neuropeptide content of each cell can be determined, and the functional effects of each signal can potentially be characterized. An elegant body of work on small circuits from crustaceans has used pharmacological and electrophysiological methods to analyze the functional effects of cotransmitters (11). However, the nature of these experimental systems typically requires that the isolated circuit be studied after dissection from the animal.

*C. elegans* provides the opportunity to use powerful genetic methods to functionally analyze cotransmission within small circuits of intact, behaving animals. Thus mutations and transgenes can manipulate neurotransmitters, neuropeptides, and their receptors, optogenetic methods can manipulate activity of presynaptic cells, and the functional consequences of all these manipulations can be read out in behavioral effects and using genetically-encoded calcium indicators to measure activity of postsynaptic cells. So far, there has been limited use of this approach to study cotransmission (18,19). One such study examined an olfactory neuron that releases glutamate to evoke a behavioral response to odor. This neuron coreleases a neuropeptide to activate a feedback loop to dampen activity of the olfactory neuron on specific timescales (18).

Here, we have applied the genetic toolbox described above to analyze the functional consequences of cotransmission by serotonin and a neuropeptide within a well-characterized small circuit of *C. elegans*. The *C. elegans* egg-laying circuit contains three neuron types that release neurotransmitters onto egg-laying muscles and each other to generate ~two minute active phases, during which rhythmic circuit activity induces egg-laying behavior, that alternate with ~20 inactive phases, during which the circuit is largely silent and no eggs are laid (20). The two serotonergic hermaphrodite specific neurons (HSNs) serve as the command neurons (21) within this circuit in that 1) worms lacking HSNs are egg-laying defective (22); 2) optogenetic activation of HSNs is sufficient to induce activity of the circuit that mimics a spontaneous active phase (23–25); 3) no other cells in the circuit have these properties (25). We show here that the HSNs use a combination of serotonin and a neuropeptide to induce the coordinated circuit activity of egg-laying active phases.

## Results

### The serotonergic HSN egg-laying neurons remain largely functional without serotonin

The small circuit that initiates egg-laying behavior is schematized in Fig 1A. The serotonergic Hermaphrodite-Specific Neurons (HSNs), along with the cholinergic Ventral Cord type C neurons (VCs), synapse onto the type 2 vulval muscles (vm2), which are electrically coupled by gap junctions to the type 1 vulval muscles (vm1) and contract with them to expel eggs (20). Loss of the HSNs results in a severe egg-laying defect: a mutation in *egl-1* causes death of the HSNs and results in animals that continue to make eggs but rarely lay them (26), resulting in the striking phenotype of adult worms distended with accumulated unlaid eggs (Fig 1B and 1C). Because addition of exogenous serotonin to worm culture media is sufficient to induce egg laying, even in worms lacking HSNs (22), it has been suggested that HSNs induce circuit activity simply by releasing serotonin, sensitizing the egg-laying muscles to activation by the acetylcholine released by other motorneurons of the circuit (25,27).

**Fig 1.**
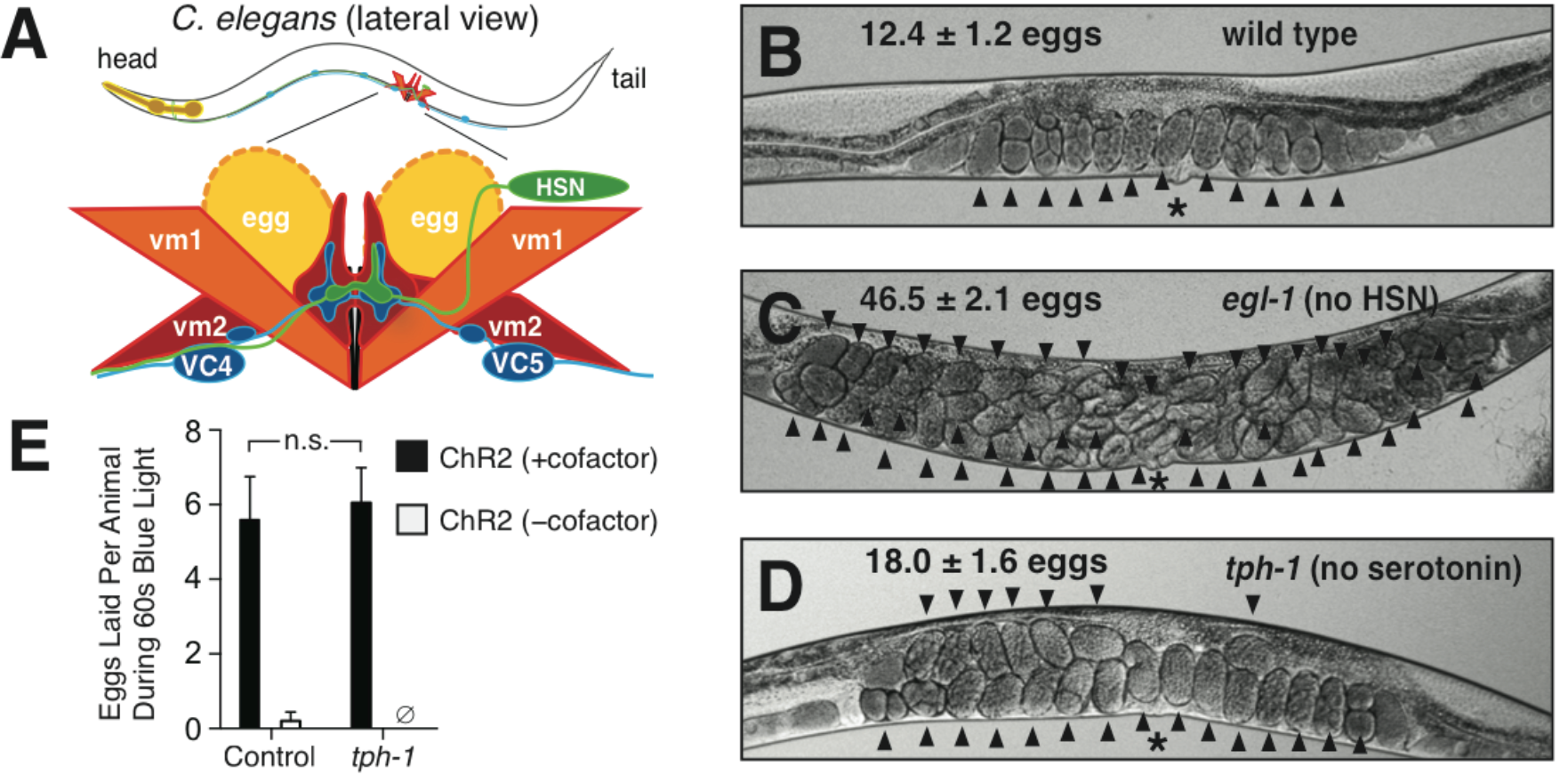
Serotonin is not required for the HSN to stimulate egg laying. **A)** Schematic of the *C. elegans* egg-laying circuit, adapted from (25). HSN and VC motorneurons synapse onto vm2 vulval muscles, which along with vm1 muscles contract to open the vulva and release eggs. Only the left HSN and vm cells are shown – equivalent cells are also found on the right side of the animal. The uv1 neuroendocrine cells that inhibit the circuit are not shown. **B-D)** Images of representative animals of the indicated genotypes, showing the average number of unlaid eggs +/− 95% confidence intervals, n=30. Arrowheads indicate individual unlaid eggs. Asterisks indicate the location of the vulva. **E)** Average number of eggs laid during 60 seconds of blue light exposure by animals expressing ChR2 in the HSNs. Both control and *tph-1* animals also have *lite-1(ce314)* mutations that eliminates a locomotion response to blue light (77). Black bars indicate animals that were grown for a generation in the presence of ChR2’s required cofactor all-trans retinal (ATR). White bars indicate negative control animals grown in the absence of ATR. Error bars, 95% confidence intervals, n=20, n.s., no statistically significant difference, ∅, no egg laying observed.

Contrary to this model, we found that animals lacking serotonin (Fig 1D) had only mild egg-laying defects. The *tph-1* gene encodes the serotonin biosynthetic enzyme tryptophan hydroxylase, and animals with a *tph-1* null mutation have no serotonin detectable by anti-serotonin antibodies or HPLC analysis (28,29). *tph-1* mutant animals had only mild egg-laying defects (~18 unlaid eggs), appearing more similar to wild type (~12 unlaid eggs) than they did to *egl-1* mutants lacking HSNs (~47 unlaid eggs; Fig 1B-1D). This result, along with previous pharmacological, genetic, and behavioral studies of the function of serotonin in egg laying (30), are consistent with the idea that serotonin release can only partially explain how the HSNs initiate egg laying.

To determine definitively if serotonin is required for the HSNs to stimulate egg laying, we optogenetically stimulated the HSNs of animals either wild-type for *tph-1* or deleted for the *tph-1* gene (Fig 1E). Animals with channelrhodopsin (ChR2) expressed in the HSNs and that are wild-type for *tph-1* have been shown previously to lay eggs within a few seconds of exposure to blue light, but only if the required ChR2 cofactor all-trans retinal (ATR) is supplied to the worms (23,24). We found that upon optogenetic activation of HSNs, *tph-1* mutant animals laid a number of eggs statistically indistinguishable from the number laid by control animals wild-type for *tph-1* (Fig 1E). Thus the HSNs do not require serotonin to stimulate egg-laying behavior.

### The neuropeptide gene *nlp-3* stimulates egg laying, and loss of both *nlp-3* and serotonin together severely reduces egg laying

Our results, along with previous studies (30–33), lead to the hypothesis that the HSN releases a cotransmitter that allows the HSNs to stimulate egg laying without serotonin. We thought that this cotransmitter could be one or more neuropeptides encoded by the five neuropeptide genes previously shown to be expressed in the HSNs (34–36). To test this idea we created five types of transgenic worm strains, each overexpressing one of these neuropeptide genes. Each strain we created carried an extrachromosomal transgene containing multiple copies of a ~45 kb *C. elegans* genomic DNA clone containing one neuropeptide gene. Overexpressing a neuropeptide gene can result in a gain-of-function phenotype caused by an increase in the normal signaling effects of the encoded neuropeptides (19,37). For a neuropeptide that induces egg laying, we expected overexpression to cause an increase in the frequency of egg-laying behavior.

We found that overexpressing the *nlp-3* neuropeptide gene resulted in a dramatic increase in the frequency of egg-laying behavior. An increased rate of egg-laying behavior results in an increased number of eggs being laid at early stages of development, since the eggs have little time to develop inside the mother before they are laid (38). More than 80% of the eggs laid by worms carrying the high-copy *nlp-3* transgene were laid at early stages of development, compared to about 5% for control animals not overexpressing any neuropeptide (Fig 2A). We saw no such phenotype for worms overexpressing any of the other four neuropeptide genes (Fig 2A).

**Fig 2.**
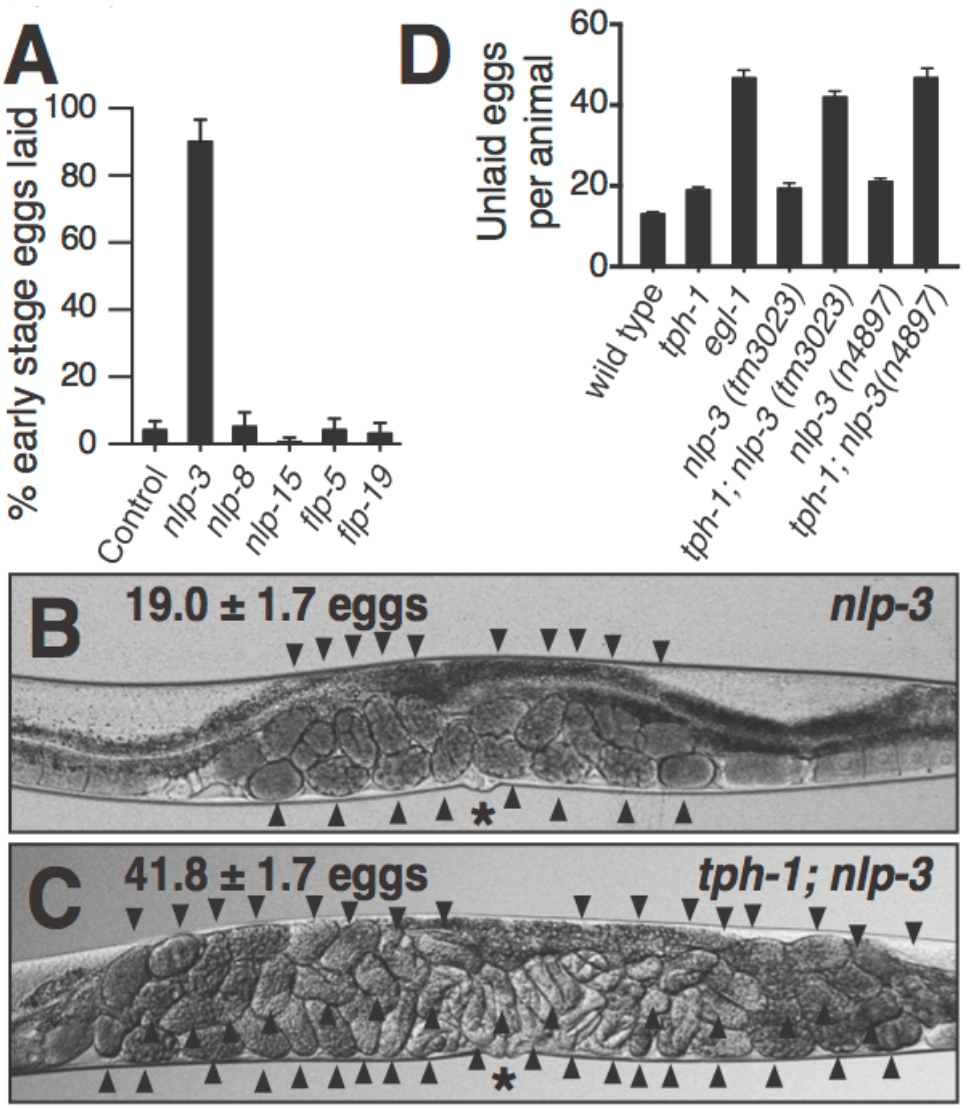
The neuropeptide gene *nlp-3*, together with serotonin, stimulates egg laying. **A)** Overexpression of *nlp-3*, but not of four other neuropeptide genes, increased the rate of egg-laying behavior. Genomic clones for each neuropeptide gene, or the coinjection marker alone (control), were injected into *C. elegans* to generate high-copy extrachromosomal transgenes. For each gene, 250 freshly laid eggs (50 from each of five independent transgenic lines) were examined and the percent laid at early stages of development (eight cells or fewer) was determined. Error bars, 95% confidence intervals. **B-C)** Representative images of *nlp-3* and *tph-1*; *nlp-3* animals showing the average number of unlaid eggs (n=30). **D)** Histogram of the average unlaid eggs for the strains indicated. Two independent deletion alleles of *nlp-3* were used. n≥30 for each strain; error bars, 95% confidence intervals. *tph-1* and the two independent *nlp-3* mutants accumulated significantly more eggs than did the wild type, and significantly fewer than did the *tph-1*; *nlp-3* double mutant strains (p≤0.05, Student’s t-test). *egl-1* mutants were not significantly different from *tph-1*; *nlp-3(n4897)*.

We obtained *nlp-3* null mutant animals in which the *nlp-3* gene is deleted. Unlike the *egl-1* mutants lacking HSNs that accumulate ~47 unlaid eggs (Fig 1C), *nlp-3* null mutants accumulated only ~19 unlaid eggs (Fig 2B), and thus were more similar to the wild type (Fig 1B) or *tph-1* mutant worms lacking serotonin (Fig 1D). However, when we made *tph-1*; *nlp-3* double mutant so that the HSNs lacked both serotonin and NLP-3 neuropeptides, the adult animals were distended with ~42 unlaid eggs and thus showed a severe egg-laying defect similar to that of *egl-1* animals. We obtained a second, independent deletion mutant for *nlp-3* and observed the same mild defect in the single mutant and the same severe defect in the double mutant with *tph-1* (Fig 2D).

### The HSN egg-laying neurons use both serotonin and NLP-3 neuropeptides to stimulate egg laying

The above experiments demonstrated that serotonin and NLP-3 stimulate egg laying but did not examine if they do so by being released from the HSN neurons. Previous studies demonstrated that HSNs contain serotonin, and it was inferred from indirect evidence that HSNs can release serotonin to stimulate egg laying (22,39,40), although our results presented in Fig 1 show that HSNs do not require serotonin to stimulate egg laying. We used the *nlp-3* promoter to drive GFP expression and saw, as previously reported (41), that *nlp-3* is expressed in the HSN neurons but no other cells of the egg-laying circuit (Fig 3A), consistent with the hypothesis that NLP-3 neuropeptides are released from HSNs to stimulate egg laying.

**Fig 3.**
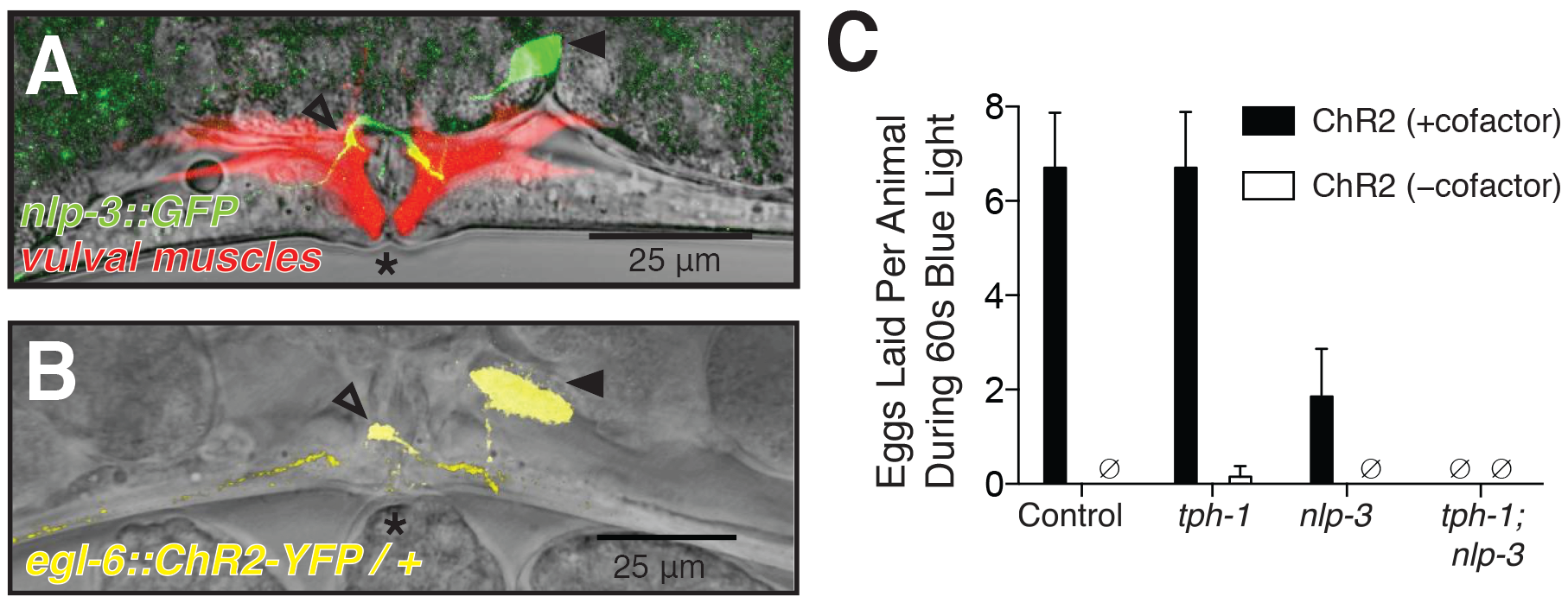
The HSNs require the *nlp-3* neuropeptide gene and *tph-1* to stimulate egg laying. **A)** *nlp-3* is expressed specifically in the HSNs. Vulval region of an adult animal carrying an nlp-3::GFP transgene, and a second transgene that expresses mCherry in the vulval muscles from the *unc-103e* promoter (46). **B)** The *egl-6p:*.ChR2-YFP transgene is expressed specifically in the HSN. In A and B, asterisks indicate the vulva. Filled arrowheads the HSN cell body, and open arrowheads the HSN synapse onto the vulval muscles. **C)** Average number of eggs laid during 60 seconds of blue light exposure by animals expressing ChR2-YFP in the HSNs, and carrying the indicated null mutations in *tph-1* and/or *nlp-3*. Control, animals wild type for *tph-1* and *nlp-3*. Black bars, animals grown in the presence of ChR2’s required cofactor all-trans retinal (ATR). White bars, animals grown in the absence of ATR. All animals in this experiment were homozygous for a *lite-1* mutation that abolished an endogenous *C. elegans* response to blue light (78). Error bars, 95% confidence intervals, n=20. ∅, no egg laying was observed.

To directly test what combination of transmitters the HSNs use to stimulate egg laying, we generated animals that express ChR-2::YFP in the HSN neurons (Fig 3B) and that were wild-type for *tph-1* and *nlp-3* (controls), or that carried null mutations in *tph-1*, *nlp-3*, or both. We then tested whether optogenetic stimulation of the HSNs could induce egg laying. Both the control animals and null mutants for *tph-1* laid eggs readily upon ChR2 activation, with no statistically significant differences in the number of eggs laid (Fig 3C), or in several other measures of the egg-laying behavior induced (e.g. time to first egg laid, time to last egg laid, Fig S2A–S2C). However, whereas all wild-type and *tph-1* animals tested laid eggs upon ChR2 activation, 7/20 *nlp-3* mutant animals failed to lay any eggs, and the 13/20 that did lay eggs laid fewer on average than did the wild-type or *tph-1* animals. No eggs were laid by any *tph-1*; *nlp-3* double mutant animals (Fig 3C). Therefore, we conclude that the HSN neurons release both serotonin and NLP-3 peptides to stimulate egg laying, either signal alone is sufficient to stimulate at least some egg laying, and when lacking both signals the HSNs have no detectable ability to stimulate the behavior.

### Serotonin and NLP-3 can each stimulate egg laying in the absence of the other

To further investigate the relationship between serotonin and NLP-3 in activating egg-laying behavior, we performed additional experiments to test if either of these transmitters is required to allow the other to stimulate egg laying. For serotonin stimulation of egg laying, we used a standard assay (22) in which worms were placed in microtiter well containing plain buffer or buffer containing serotonin, and the number of eggs laid in 60 minutes was counted. We saw, as observed previously (39,42–44), that exogenous serotonin stimulates egg laying in wild-type animals, but not in animals deleted for the serotonin receptor gene *ser-1* (Fig 4A). Null mutants for *nlp-3* were stimulated by serotonin to lay eggs at the same rate as were the wild-type controls, demonstrating that NLP-3 is not required for serotonin to stimulate egg laying.

**Fig 4.**
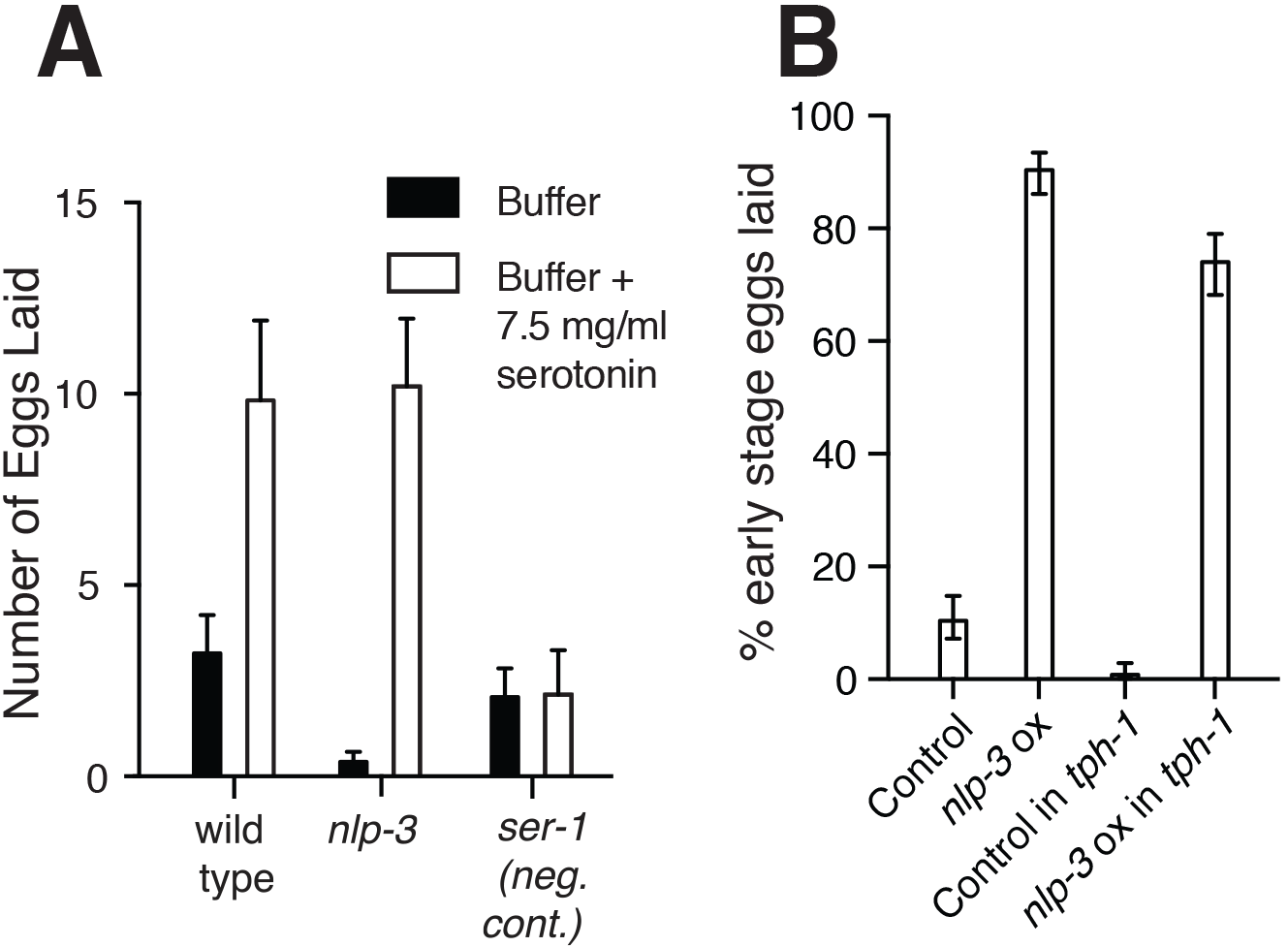
Serotonin and NLP-3 neuropeptides can stimulate egg laying in the absence of each other. **A)** Exogenous serotonin stimulates egg laying in wild type and *nlp-3* animals. The number of eggs laid by 10 animals over 30 minutes in plain buffer or buffer plus serotonin was measured, averaging 10 replicates per genotype. The *ser-1* serotonin receptor null mutant is the negative control. **B)** *nlp-3* overexpression stimulates egg laying even in the absence of serotonin. Animals wild-type for *tph-1* or *tph-1* null mutants were injected with marker DNA alone (control), or *nlp-3* genomic DNA plus marker DNA to overexpress *nlp-3* (*nlp-3* ox). In each case, five independent transgenic lines were produced, and 50 freshly laid eggs per line (250 eggs total per condition) were examined to determine their developmental stages. Error bars, 95% confidence intervals.

We used a converse experiment to test if NLP-3 could stimulate egg-laying in the absence of serotonin. We generated *C. elegans* transgenes that overexpressed *nlp-3* by containing multiple copies of *nlp-3* genomic DNA, and control transgenes that did not overexpress *nlp-3*. In a strain background wild-type for *tph-1*, we observed (Fig 4B), as we had seen previously in an analogous experiment (Fig 2A), that overexpression of *nlp-3* resulted in hyperactive egg laying as evidenced by a high percentage of early-stage eggs laid. When we carried out this same experiment in a *tph-1* null mutant, *nlp-3* overexpression also resulted in hyperactive egg laying, albeit at a modestly reduced level (Fig 4B). Thus serotonin is not required to allow *nlp-3* overexpression to induce egg laying.

### Serotonin and NLP-3 together cause the HSN postsynaptic targets, the vm2 muscle cells, to contract coordinately with other egg-laying muscles

To understand the functional effects of the HSN cotransmitters, we recorded Ca^2+^ activity of the vulval muscle cells, which are postsynaptic targets of the HSNs, in animals that were wild-type, lacked serotonin, lacked NLP-3, or lacked both. We thus coexpressed the Ca^2+^-sensitive green fluorescent protein GCaMP5 and the Ca^2+^-insensitive red fluorescent protein mCherry in the vm2 muscle cells, the direct postsynaptic targets of the HSNs, and also in the vm1 muscle cells, which are gap-junctioned to vm2 and have been thought to contract with vm2 to expel eggs (45). Using methods we previously developed (25,46,47), we carried out ratiometric fluorescence imaging of intact animals to measure Ca^2+^ transients under conditions that allow egg-laying behavior to proceed as it does in standard lab culture, such that in wild-type animals, ~2 minute egg-laying active phases occur about every 20 minutes. Each animal was recorded for one hour. Fig 5A shows traces of Ca^2+^ transients recorded for the entire ensemble of vm1 and vm2 cells together for three animals of each of the following genotypes: wild-type, *tph-1* and *nlp-3* single mutants, the *tph-1*;*nlp-3* double mutant, and *egl-1* animals lacking HSNs.

**Fig 5.**
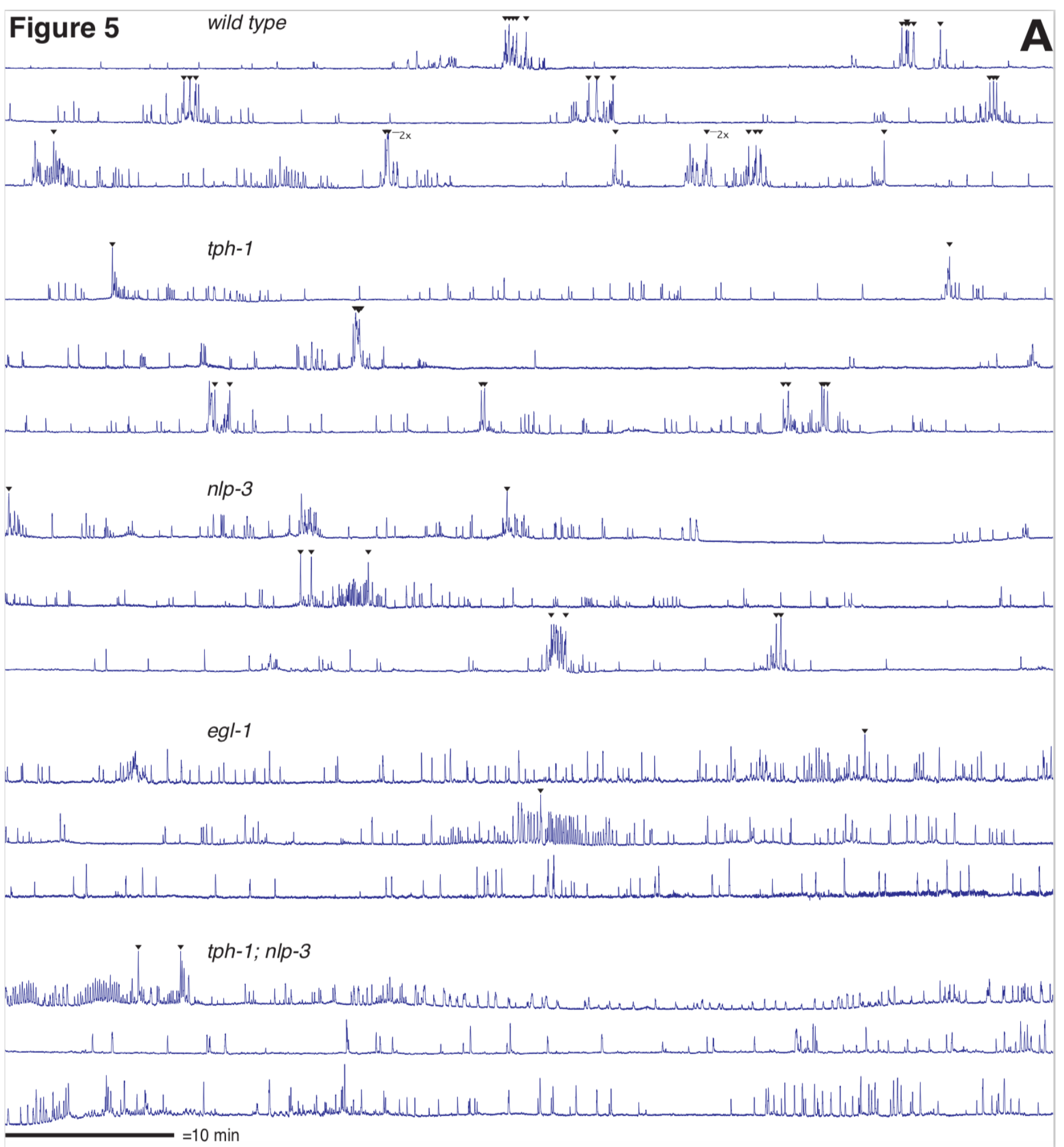
Vulval muscle activity, but not egg release, occurs frequently in mutants lacking serotonin, NLP-3, or both. **A)** Graphs of Ca^2+^ transients showing ΔR/R of GCaMP5/mCherry signal recorded over 1 hour for three different animals per genotype. ▼indicates a calcium transient associated with an egg-laying event. “2x” indicates that two eggs were laid nearly simultaneously during the same calcium transient. Scale bar, 10 minutes. Vertical scales have been normalized to depict comparable peak heights in all animals shown. **B)** Still frames from ratiometric recordings. The mCherry channel is rendered in blue. The mCherry channel is rendered in green, with a scale of intensity ranging from transparent, low intensity to greener high intensity. Schematics are shown below the images to distinguish the muscle types and indicate where activity appears to occur. **C)** A graph of the number and proportion of calcium transients occurring in the vm1s only compared to those appearing to occur in both vm1 and vm2 for each genotype. Error bars show 95% confidence intervals for the proportion that would result from an infinite number of observations.

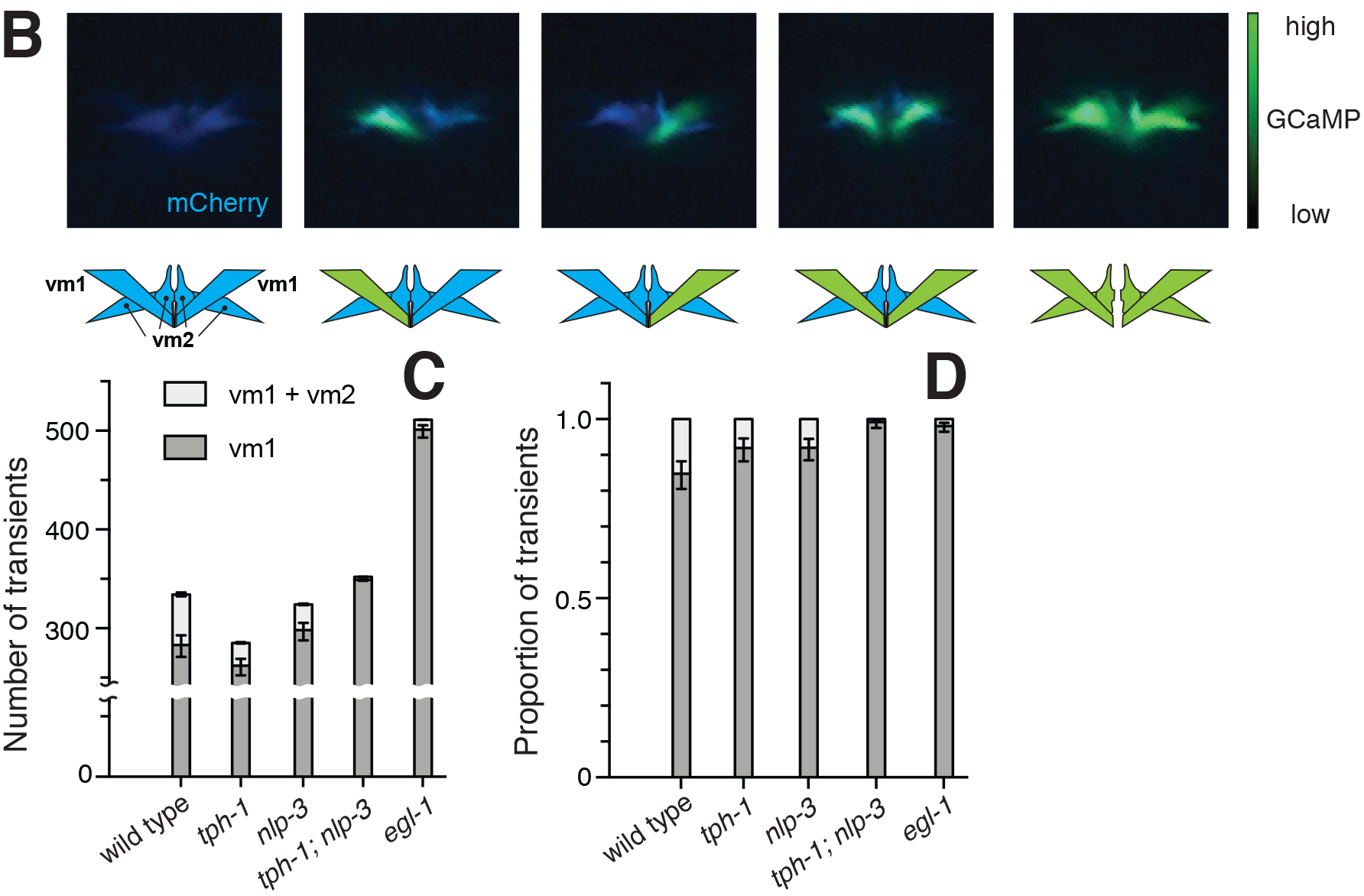

We observed frequent vulval muscle activity in all genotypes, with each genotype showing hundreds of Ca^2+^ transients over the three hours recorded. However, in the wild-type less than 10% of the vulval muscle Ca^2^+ transients resulted in egg release, and even fewer successful egg-laying events occurred in *tph-1* mutants lacking serotonin (*tph-1*) or in *nlp-3* mutants (30 eggs released over three hours for the wild type, compared to 15 for *tph-1* and 9 for *nlp-3)*. Activity in animals lacking both serotonin and NLP-3 neuropeptides *(tph-1; nlp-3)* or lacking HSNs (*egl-1*) was actually more frequent than in the wild type, but very rarely produced successful egg release (each genotype released just two eggs in the three hours recorded).

To identify the differences between vulval muscle contractions that did or did not release eggs, we adjusted how we collected images during Ca^2+^ recordings. Previously-published Ca^2+^ imaging of the vulval muscles used images focused at the center of the group of two vm1 and two vm2 muscles found on either the left or right side of the animal, and the resulting images showed Ca^2+^ activity that was usually focused at the most dorsal tip of this group of muscles, but that could not be assigned to individual muscle cells (25,46,47). By focusing more laterally on either the left or right set of vulval muscles, we could more clearly resolve individual vm1 and vm2 cells and determine which of the four muscle cells within the set were active during any given Ca2+ transient detected (Fig 5B). All of the data presented in Fig 5 and Fig S3 results from use of this more lateral focus.

The large majority of the muscle activity we observed in every genotype examined occurred exclusively in one both of the vm1s imaged, with no concurrent activity detected in the vm2s (Fig 5C). In the wild-type, 12% of Ca2+ transients involved both vm1 and both vm2 cells imaged, and we refer to such events as "coordinated". We never observed an event in any genotype in which a Ca^2+^ transient occurred exclusively in vm2 cell(s) without accompanying activity in vm1 cell(s). In the wild-type, coordinated vulval muscle contractions occurred exclusively within active phases, the ~2 minute intervals during which eggs were laid and that contained frequent vulval muscle transients (Fig 5A). All 30 egg release events observed in the wild type occurred during one of the 51 coordinated vulval muscle contractions we saw during the three hours of recordings analyzed. Thus it appears that coordinated contraction of all the vulval muscle cells is necessary for efficient egg release.

Mutants lacking serotonin, NLP-3, or both continued to show vm1 Ca2+ transients at a rate similar to or even greater than seen in the wild-type, but a decreased portion of these events were accompanied by vm2 Ca2+ transients to produce coordinated events (Fig 5C). This decrease in coordinated events was modest in *tph-1* and *nlp-3* single mutants, but severe in the *tph-1*; *nlp-3* double mutant and in *egl-1* animals lacking HSNs. In the mutants, as in the wild-type, egg release occurred almost only during coordinated events that included both vm1 and vm2 activity (Fig S3): we only observed one exceptional egg-release event that occurred in an *egl-1* animal during a vm1-only contraction. Thus the loss of vm2 activity in the mutants correlated strongly with the loss of successful egg laying. Thus, while vm1 calcium activity occurs in each genotype we observed, only wild-type animals, in which the HSN neurons were able to release both serotonin and NLP-3 neuropeptides onto vm2 cells, were able to frequently trigger coordinated activity in both vm1 and vm2 and efficiently lay eggs.

## Discussion

### HSN command neurons release serotonin and NLP-3 neuropeptides to activate and coordinate activity of the egg-laying circuit

Our results show that serotonin and NLP-3 released by the HSNs induces activity of the vm2 muscle cells and coordinates their activity with that of the vm1 muscles to productively release eggs. In wild-type animals, the egg-laying circuit alternates between ~20 minute inactive states during which no eggs are laid, and ~2-3 minute active states during which a few eggs are laid. Previous work (21,42,43) as well as results in this study show that there are occasional vm “twitch” Ca^2+^ transients during the inactive state that do not release eggs, while there are robust, rhythmic vm transients during the active state, some of which release eggs. There are a total of four vm1 and four vm2 cells, with Fig. 1 diagramming just the two of each type on the left side of the animal. Gap junctions are present among the vm1 and vm2 cells found either anterior or posterior to the vulva, but there are no gap junctions between the anterior and posterior sets of vm cells (45). A previous study (48) showed that egg-laying events always coincide with Ca^2+^ transients that occur simultaneously in both anterior and posterior vm cells, while animals lacking HSNs fail to show such anterior/posterior vm coordination, potentially explaining the lack of efficient egg laying in animals lacking HSNs. It was assumed in such previous studies that the gap junctions between vm1 and vm2 muscle cells anterior or posterior on either side of the vulva would efficiently electrically couple these cells so that they would contract as unit. In this study, we increased the spatial resolution of our Ca^2+^ imaging and found that this assumption was incorrect. We observed that the vm twitch Ca^2+^ transients seen in the inactive state occur in some or all of the vm1 cells but are not detected in the vm2 cells. During the active state in wild-type animals, a subset of the rhythmic vm transients seen happen simultaneously in all vm1 and vm2 cells, and it is in turn a subset these “coordinated” events that result in egg release. Animals lacking the HSNs or the HSN-released signals continue to have vm1 Ca^2^+ transients, but show a profound loss of vm2 Ca^2+^ transients. Thus the HSNs and their signals are not necessary for vm1 activity, which apparently are stimulated by other source(s), but the HSNs are critical for stimulating vm2 activity so that all vm cells can contract coordinately to productively release eggs.

The anatomy of the egg-laying circuit helps explain how the vm1 and vm2 muscle cells are activated. Figure 6 diagrams the synapses and gap junctions among cells in the circuit as determined by serial section reconstruction of electron micrographs (EM) (45). The HSNs synapse onto the vm2 muscles, as do the cholinergic VC neurons. In past studies, HSN and VC were considered to be the primary motor neurons that stimulate the vm muscles, but our results show that the vm1 cells, which are not synaptic targets of HSN or VC neurons, are activated independently of vm2. What cells and signals then excite the vm1 muscles?

**Fig 6.**
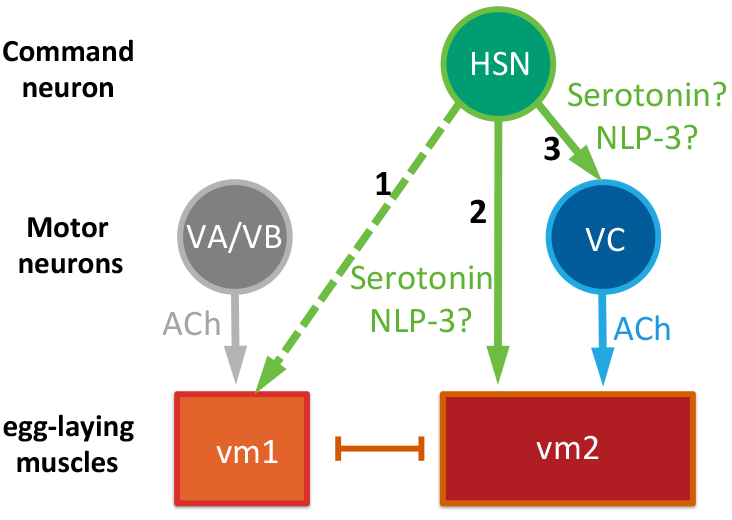
Model depicting signaling events that activate the egg-laying circuit. Solid arrows, synaptic signaling; dashed arrow, extrasynaptic signaling; and bar, gap junctions. vm1 and vm2 cells are known to express multiple serotonin receptor isoforms that each mediate activation of egg laying. The signaling of serotonin onto VC neurons (arrow 3) remains hypothetical since no serotonin receptors have yet been described as expressed on these neurons. The possible direct signaling of NLP-3 depicted onto vm1 (arrow 2), vm2 (arrow 2), or and/or VC (arrow 3) are also hypothetical: NLP-3 receptors have yet to be identified, and it thus remains unknown which cell(s) of the circuit express them.

We hypothesize that the VA and VB motor neurons are the source of excitation of the vm1 muscles, and also the central pattern generators that produce rhythmic activity of the egg-laying muscles. EM reconstruction (45) shows that the vm1 cells receive synapses from the cholinergic VA7 and VB6 motor neurons, although these synapses have been disregarded in the past because they are much smaller than are the VC and HSN synapses onto vm2. VA and VB are ventral cord motor neuron classes that synaptically release acetylcholine onto the body wall muscles to coordinate the body bends of locomotion. Rhythmic behaviors such as locomotion generally require central pattern generator (CPG) neurons that are the source of their rhythmicity (49). VA and VB are rhythmically active and serve as CPGs that produce the pattern of body bends during locomotion (50). During the active state, egg laying is a repetitive behavior (27), and there must be a connection between the CPG for locomotion and the egg-laying circuit because repetitive vm activity is phased with the body bends of locomotion (25,46). Cholinergic activation of vm1 by VA/VB neurons would neatly explain both the source of vm1 activity and the source of the rhythmicity of vm activity. Previous work showed that vm muscles are apparently stimulated at a specific phase of every body bend, even during the inactive state, since mutations in specific K^+^ channels that normally reduce excitability of the vm cells result in increased rhythmic vm contractions, phased with body bends, during both the inactive and active egg-laying states. VA/VB neurons are active at a specific phase of each body bend (50) regardless of whether animals are in the inactive or active state of the egg-laying system, and thus their pattern of activity exactly matches the pattern of activity inferred for the previously unknown source of vm activity.

We found that HSN neurons release serotonin and NLP-3 to initiate the egg-laying active state. Prevoius work showed that optogenetic activation of HSNs is sufficient to stimulate egg laying (24), and induces a pattern of activity in the VC neurons and in vm muscles reminiscent of activity seen during spontaneous active states (25). In this work, we found that optogenetic stimulation of HSNs lacking either serotonin or NLP-3 can still induce egg laying, but loss of both eliminates any egg laying response. Therefore, it appears that serotonin and NPL-3 together allow the HSNs to induce the active state of the egg-laying circuit. Examining spontaneous activity of the egg-laying circuit (i.e. without optogenetic activation of HSNs), we see that elimination of serotonin, NLP-3, both, or even killing the HSNs with an *egl-1* mutation does not eliminate egg-laying circuit activity. Indeed, vm Ca^2+^ activity remains high in these animals, and even includes activity in clusters, a property of the spontaneous active state seen in the wild type. However, the vm activity in animals lacking the HSN or its signals is uncoordinated, with Ca^2+^ transients mainly seen in vm1 cells only. Coordinated events with simultaneous Ca^2+^ transients in all vm cells that result in egg release occur on occasion, but these tend to be isolated events, as opposed to the groups of several coordinated egg-laying contractions that tend to occur within a 2-3 minute active phase in the wild type. The vm1-only activity seen in mutants lacking the HSN or its signals could arise simply from direct release of acetylcholine onto vm1 cells by the VA/VB neurons. We also know from our previous work that it depends on an unknown signal released when the uterus contains unlaid eggs (25): the bloating of the uterus with excess unlaid eggs in animals lacking the HSNs or its signals may increase the uterus signal to result in the high levels of vm1-only activity seen in such animals.

How do HSNs use serotonin to produce vm2 contractions and the egg-laying active phase? The HSN and VC neurons make synapses onto muscle arms that project from all vm2 muscle cells, including those both anterior and posterior to the vulva. This anatomy ideally positions HSN and VC to stimulate vm2 contractions, and to do so such that both the anterior and posterior sets of vm cells contract simultaneously to productively release eggs. Previous studies showed that serotonin stimulates egg laying via the SER-1, SER-7, and SER-5 G protein coupled receptors (GPCRs), with potentially additional help from the MOD-1 serotonin-gated ion channel (51). Promoter::GFP transgenes for the *ser-1*, *ser-7*, and *ser-5* receptor genes show expression in the vm cells (32,42,44,51,52). The transgenic animals carrying these GFP reporters have not been examined carefully to determine which receptors are expressed in vm1 versus vm2 cells, but the published images suggest that both vm1 and vm2 express one or more of these GPCRs. No expression of *mod-1*::GFP has yet been seen in the egg-laying circuit (53). Thus, the anatomy of the egg-laying circuit and serotonin receptor expression patterns suggest that HSN-release serotonin is released at synapses directly onto vm2 cells to activate these muscles via one or more GPCRs, and additionally activates vm1 cells extrasynaptically via one or more GPCRs. As a neuromodulator, serotonin could increase the excitability of vm1 cells and thus their depolarization upon release of acetylcholine onto vm1 by the VA/VB neurons. Serotonin could similarly act on vm2 cells to increase their response to acetylcholine released onto vm2 by the VC neurons, and/or their ability to respond to depolarization via gap junctions from vm1 cells.

How do HSNs use NLP-3 neuropeptides to produce vm2 contractions and the egg-laying active phase? Hypotheses for the action of NLP-3 are more speculative than are those for the action of serotonin, as NLP-3 receptors have not yet been identified, and we thus do not yet know which cells express these receptors. NLP-3 might act on the vm cells just as does serotonin, so that these two signals would activate the egg-laying circuit in the same manner. However, a more interesting hypothesis is that NLP-3 is the signal that the HSNs use to activate the VC neurons. Previous studies show that the VC neurons are essentially silent during the egg-laying inactive state, but become rhythmically active during the egg-laying active state, and that optogenetically activating the HSN neurons is sufficient to induce such activity of the VC neurons (52). Although the expression pattern of every known serotonin receptor in *C. elegans* has been described, none have so far been seen expressed on the VC neurons. Therefore, an attractive model is that NLP-3 is the HSN-released signal that acts on the VC neurons to induce their activity. Active VC neurons would then release acetylcholine directly onto the vm2 muscles. In this way, HSN-released serotonin and NLP-3 would each have an independent mechanism for inducing vm2 activity. The possible targets of signaling by serotonin and NLP-3 outlined in the model described above are depicted in Figure 6. This model is consistent with our experimental observations that HSN-released serotonin and NLP-3 can each independently induce egg laying.

### Co-release of small molecule neurotransmitters and neuropeptides is a widespread phenomenon

The apparent co-release of serotonin and NLP-3 from the HSN neurons is just one instance of the broad but poorly-studied phenomenon of co-transmission by small-molecule neurotransmitters and neuropeptides. Neurons typically release one (or more rarely more than one) small molecule neurotransmitter from small synaptic vesicles (SSVs) (54,55), and release neuropeptides from large dense-core vesicles (LDCVs) (11,56). Certain small-molecule neurotransmitters, including serotonin, can also be found in LDCVs (54,55). SSVs and LCDVs can be localized in different parts of the cell and released by different mechanisms (55,57).

Most neurons release both small-molecule neurotransmitters and neuropeptides. This issue has been analyzed in greatest detail within the *C. elegans* nervous system. There are 118 neuron types in *C. elegans* hermaphrodites, and 107 of them express one or more of the seven known small molecule neurotransmitters found in this organism (58). At least 95 *C. elegans* neuropeptide genes have been described, including 23 FLP genes encoding FMRFamide-related peptides, 32 NLP genes encoding neuropeptide-like proteins, and 40 INS genes encoding insulin-like peptides. Promoter::GFP fusion transgenes have been generated for all 95 of these neuropeptide genes to analyze their expression patterns. The individual neurons expressing each FLP gene were identified, and >50% of *C. elegans* neurons express at least one FLP peptide gene (35). The individual cells expressing each NLP and INS gene have not yet been identified, but images of the expression patterns show that the large majority of these peptide genes are expressed complex subsets of neurons (34,59). Thus, we can infer that the typical neuron in *C. elegans* releases one small-molecule neurotransmitter, and one or more type of neuropeptide. Similarly, the presence of both small molecule neurotransmitters and neuropeptides within the same individual neurons is widespread in both *Drosophila* (60,61) and in mammals (62).

The functional consequences of a neuron releasing two different types of signaling molecules have been difficult to study with precision in the complex circuits of the mammalian brain, but this issue has been the focus of many studies of small neural circuits in invertebrate model organisms (11,63). In such small circuits, individual presynaptic neurons that co-release a small molecule neurotransmitter and neuropeptides can be identified, and the functional effects of each signal can be measured by bath application of neurotransmitter agonists/antagonists and/or neuropeptides, followed by measurements of circuit activity using electrophysiological methods. Such work has led to a rich set of findings, and many different schemes for the use of co-transmission within circuits (11,64). However, the limitations of these studies include that bath application of signaling molecules does not always mimic the effects of their release from neurons (11). Further, the electrophysiological recordings used require dissecting neural circuits out of the animal, replacing their extracellular fluid with an artificial solution, eliminating the movements that motor circuits normally induce and that provide proprioceptive feedback to these circuits, and impaling the recorded neurons with electrodes that dialyze their intracellular fluid, all of which may affect circuit function. The genetic approaches for analyzing cotransmission described in this work provides a useful complement to electrophysiological studies, as they permit us to manipulate endogenous signaling molecules with mutations and transgenes, to record circuit activity using genetically-encoded calcium indicators, and to manipulate neural activity using optogenetics, all within intact, freely-behaving animals. We are aware of just one previous study that focused on co-transmission using this combination of genetic approaches (18). In this pioneering study, an odor was shown to cause a *C. elegans* sensory neuron to release glutamate to act via ionotropic receptors on specific interneurons that further regulate a complex and incompletely understood motor circuit to evoke a behavioral response to the odor. The same sensory neuron also releases a neuropeptide that acts via a G protein coupled receptor on a different interneuron to cause it to in turn release a second neuropeptide back onto the sensory neuron, limiting activity of the sensory neuron and the timescale of the behavioral response to the odor.

Our studies of co-transmission focus on the *C. elegans* egg-laying circuit because its anatomical simplicity holds the promise that all the cells and signaling events that control this circuit can be defined, something that has not yet been accomplished for any neural circuit. We discovered that serotonin and NLP-3 peptides released from the HSN command neurons have parallel and partially redundant effects to activate coordinated, rhythmic contraction of the egg-laying muscles. This finding may be analogous to results of some previous studies of co-transmission, in which the two co-released signals act convergently to increase activity the same target cells. The most relevant such example is in the mammalian brain respiratory circuit, where co-release of serotonin and the neuropeptide Substance P have parallel effects promoting rhythmic circuit activity (10). It will be interesting to determine just how mechanistically analogous these two cases of serotonin/neuropeptide co-transmission actually are, and whether the action of serotonin within the *C. elegans* egg-laying circuit will provide a model for the detailed workings of serotonin within neural circuits of the human brain.

## Methods

### *C. elegans* strains

*C. elegans* strains were cultured at 20°C on NGM agar plates with *E. coli* strain OP50 as a food source (66). All strains were derived from the Bristol N2 wild-type strain. Genetic crosses and generation of transgenic strains were by standard methods (67,68). A list of strains, mutants, and transgenes used in this study can be found in Table 1.

Gene deletion strains were for *nlp-3 (69)*, *tph-1 (70)*, and *ser-1* (42)were outcrossed four to ten times to the wild-type strain, as was the strain carrying an additional, previously unpublished *nlp-3* deletion allele, *tm3023*, which we obtained from the Japanese National Bioresource Project. It carries a 354 bp deletion that removes sequences flanked by the sequences GTCTGGACGGAAAGATCGTT…CGTGAGACTAGAAGTCCAC. Each gene deletion used removes a portion or all of the promoter and/or coding sequences of the corresponding gene such that no functional gene product is expected. The genotypes for all strains constructed using these deletions were verified by agarose gel analysis of PCR amplification products from the corresponding genes.

**Table 1.** Strains used in this study.

| Strain | Feature | Genotype | Figures |
| --- | --- | --- | --- |
| N2 | Bristol strain | Wild type | 1,2 |
| MT2059 | Lacks HSN neurons | *egl-1(n986dm)* V | 1,2 |
| MT15434 | Lacks serotonin | *tph-1(mg280)* II | 1,2 |
| LX1836 | *egl-6::ChR2::YFP* | *wzls30 IV; lite-1(ce314) lin-15(n765ts)* X | 1,3 |
| MT8189 | Strain for transgene production | *lin-15(n765ts)* X | 2 |
| LX1954 | *nlp-3* overexpressor | *lin-15(n765ts)* X ; *vsEx748* | 2 |
| LX1955 | *nlp-8* overexpressor | *lin-15(n765ts)* X ; *vsEx749* | 2 |
| LX1956 | *nlp-15* overexpressor | *lin-15(n765ts)* X ; *vsEx750* | 2 |
| LX1957 | *flp-5* overexpressor | *lin-15(n765ts)* X ; *vsEx751* | 2 |
| LX1981 | *flp-19* overexpressor | *lin-15(n765ts)* X ; *vsEx757* | 2 |
| LX1978 | *nlp-3* null mutant | *nlp-3(tm3023)* X | 2,4 |
| LX2366 | double mutant | *tph-1(mg280)* II; *nlp-3(tm3023)* X | 2 |
| LX2388 | *nlp-3* null mutant | *nlp-3(n4897)* X | 2 |
| LX2389 | double mutant | *tph-1(mg280)* II; *nlp-3(n4897)* X | 2 |
| LX1836 |  | *wzls30 IV; lite-1(ce314) lin-15(n765ts)* X | 3 |
| LX1832 | mate to LX1836 for Fig 3C “control” | *lite-1(ce314) lin-15(n765ts)* X | 3 |
| LX1837 |  | *tph-1(mg280)* II; *wzls30 IV; lite-1(ce314) lin-15(n765ts)* X | 3 |
| LX2335 | mate to LX1837 for Fig 3C “*tph-1*” | *tph-1(mg280)* II; *lite-1(ce314) lin-15(n765ts)* X | 3 |
| LX2367 |  | *wzls30 IV; lite-1(ce314) nlp-3(tm3023) lin-15(n765ts)* X | 3 |
| LX2364 | mate to LX2367 for Fig 3C “*nlp-3*” | *lite-1(ce314) nlp-3(tm3023) lin-15(n765ts)* X | 3 |
| LX2368 |  | *tph-1(mg280)* II; *wzls30 IV; lite-1(ce314) nlp-3(tm3023) lin-15(n765ts)* X | 3 |
| LX2365 | mate to LX2368 for Fig 3C “*tph-1; nlp-3*” | *tph-1(mg280)* II; *lite-1(ce314) nlp-3(tm3023) lin-15(n765ts)* X | 3 |
| DA1814 | Serotonin receptor 1 deletion | *ser-1(ok345)* X | 4 |
| LX2392 | “control” in Fig 4B | *lin-15(nt65ts)* X; *vsEx885* | 4 |
| LX2394 | “*nlp-3* ox” in Fig 4B | *lin-15(nt65ts)* X; *vsEx887* | 4 |
| LX2393 | “control in *tph-1*” in Fig 4B | *tph-1(mg280)* II; *lin-15(nt65ts)* X; *vsEx886* | 4 |
| LX2395 | “*nlp-3* ox in *tph-1*” in Fig 4B | *tph-1(mg280)* II; *lin-15(nt65ts)* X; *vsEx888* | 4 |

### Egg-laying behavioral assays

Quantitation of unlaid eggs in adult animals and percentage of early-stage eggs laid was done as described in (71), using adult animals 30 hours after staging as late L4 larvae.

### Optogenetic assays

HSN neurons were optogenetically activated in animals carrying the *wzIs30* transgene, which expresses a Channelrhodopsin-2::yellow fluorescent protein (ChR2::YFP) fusion in the HSN and a few other neurons unrelated to the egg-laying circuit from the *egl-6a* promoter (72,73). *wzIs30* also carries a *lin-15* marker plasmid that rescues the multivulva phenotype of *lin-15* mutant animals. All animals used in optogenetic assay were also mutant for the *lite-1* gene to eliminate an endogenous response of *C. elegans* to blue light. The *wzIs30* transgene was homozygous for the experiment shown in Fig 1E, but we noticed that the homozygous transgene caused developmental defects in the HSNs of some animals (Fig S1) that resulted in these animals being egg-laying defective. Therefore, for the experiment in Fig 1E, we examined the animals prior to optogenetic stimulation and discarded the small percentage of animals that were visibly egg-laying defective. The experiment shown in Fig 3C was carried out such that all animals were *wzIs30/+* heterozygotes, which we found had morphologically normal HSNs (Fig S1). First we constructed the strains indicated in Table 1 that were homozygous for *wzIs30* and also homozygous for the other mutations required by the experiment. We generated males of each of these strains, and mated them to corresponding strains that were genetically identical except that they lacked *wzIs30*. The cross progeny, identified by the presence of YFP-labeling, thus were heterozygous for *wzIs30* but homozygous for all other mutations used in the experiment.

ChR2 expressing strains were grown in the presence or absence of the ChR2 cofactor all-trans retinal (ATR). ATR was prepared at 100 mM in 100% ethanol and stored at 20° C. To prepare NGM plates for behavior analysis, ATR was diluted to 0.4 mM with warmed cultures of 0P50 bacteria in B Broth, and 200 ml of culture was seeded onto each 60 mm NGM plate. The plates were allowed to grow for 24 hr at 25–37 °C, after which late L4 worms were staged onto prepared plates for behavioral assays 24 hr later. To initiate an assay, the shutter was opened to initiate exposure to blue light simultaneously with a recording (Flea 3, 0.3 Megapixel, FireWire CCD camera, Point Grey Research) and shutter opening on a EL6000 metal halide light source generating 10 mW/cm of 470 ± 20 nm blue light via a EGFP filter set mounted on a Leica M165FC stereomicroscope.

### Molecular biology and transgenes

For overexpression of neuropeptide genes (Fig 2A), fosmid genomic clones including individual neuropeptide genes were selected from the *C. elegans* fosmid library (74,75). The fosmids used for four neuropeptide genes were: *nlp-3*, WRM0633dC06; *nlp-8*, WRM0614aB10; *nlp-15*, WRM066cH12; *flp-5*, WRM0622aF03. For overexpression of a fifth neuropeptide gene, *flp-19*, we instead PCR amplified genomic DNA containing the 746 bp *flp-19* coding region along with 5015 bp upstream and 746 bp downstream. Multicopy extrachromosomal transgenes were generated for each neuropeptide gene by microinjection (reference), using the fosmid or PCR product at 50 ng/μl along with the *lin-15* rescuing plasmid pL15EK at 50 ng/μl into *lin-15(n765ts)* mutant animals. Negative controls were injected with pL15EK without any neuropeptide gene. Five independent transgenic overexpressor lines were generated for each injection and Fig 2A shows data averaged from these. Table 1 lists one representative overexpressor strain for each neuropeptide gene.

For determine the effects of overexpressing *nlp-3* in animals lacking serotonin (Fig 4B), either a ~5 kb PCR product containing the *nlp-3* gene (primers used were 5’-accaagctaatcaaattttgtcaccg-3’ and 5’-gcaatacaaccaatcccttttcatctc-3’) or as a control, *E. coli* genomic DNA digested to an average size of ~5 kb, was injected at 10 ng/μl along with 50 ng/μl of the *lin-15* rescuing plasmid pL15EK into either *lin-15* or *tph-1*; *lin-15* animals, and transgenic lines were identified by rescue of the *lin-15* phenotype. Five independent transgenic lines were established for each injection, and the early stage egg assay (76) was carried out on 50 eggs per line (250 eggs total per condition tested). One representative line for each condition is listed in Table 1.

### Ratiometric Calcium Imaging

Freely-behaving animals were mounted between a glass coverslip and chunked section of an NGM plate for imaging as described (25,46,47) and recorded with a 20X Plan-Apochromat objective (0.8 NA) using a Zeiss LSM 710 Duo LIVE head set to record two channels. Recordings were collected at 20 fps at 256 × 256 pixel, 16 bit resolution, for 1 hour. The stage and focus were adjusted manually to keep the egg-laying system in view and focused during recording periods. Care was taken to find a lateral focus that included as much of the vm1s and vm2s as possible. Ratiometric analysis for Ca^2+^ recordings was performed in Volocity (version 5, PerkinElmer). A ratio channel was calculated from GCaMP5 (GFP) and mCherry fluorescence channels. Volocity was also used to identify the vulval muscles using size and intensity parameters that varied over a small range based on individual animals. Any misidentified objects were manually excluded prior to final analysis. The lowest 10% of the GCaMP5/mCherry ratio values were averaged to establish a Δ*R/R* baseline using a custom Matlab script. This script also identifies the peak of a transient based on identifying a change in prominence that was typically 0.25 ΔR/R over the preceding second, but this was adjusted based on the smoothness of the data for individual animals. With the experimenter blinded to the genotype of the animals being scored, video of each peak was observed in the ratio channel to determine whether the indicated activity was restricted to vm1 or present in both vm1 and vm2, and whether an egg was laid. We scored a transient as vm1-only if it was clear in the ratio channel that there was a difference of more than 50% of maximum activity between the vm1s and the adjacent regions where vm2 cells were located.

### Statistical Methods

Statistical analyses were performed using GraphPad Prism for Mac OS X v. 7.0a. 95% confidence intervals were determined and 1- or 2-way ANOVA with multiple comparisons were performed to determine statistical significance. For egg stage assays, we used the Wilson-Brown method for determining the 95% confidence intervals for binomial data.

## Acknowledgements

We thank Kelly Culhane for assistance with exogenous serotonin assay.

## Supporting Information Captions

**Fig S1.**
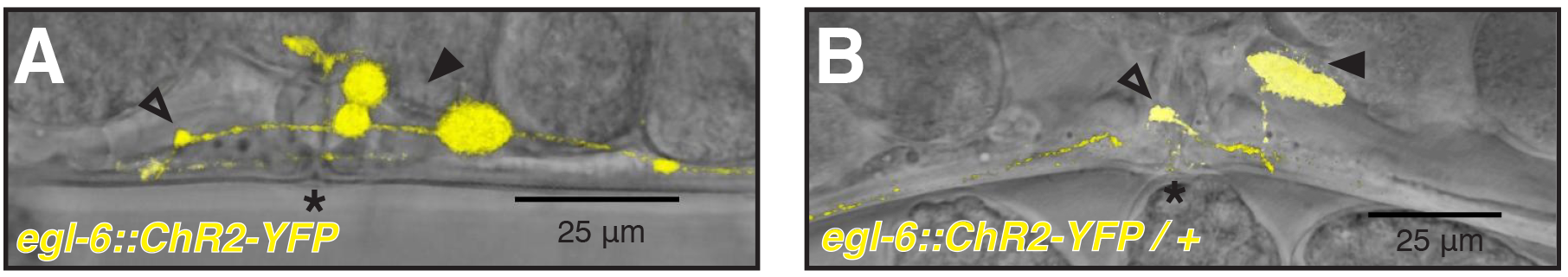
Animals homozygous for the *egl-6*::ChR2-YFP transgene have visibly defective HSNs. **A-B)** Vulval region of an adult homozygote for the *egl-6p*::ChR2::YFP transgene. Abnormal HSN morphology can be seen by comparing to normal HSN morphology in Fig. S1B. Asterisk, location of the vulva. B) Vulval region of an adult heterozygote for the *egl-6p*::ChR2::YFP transgene. This is the same image seen in Fig 3B, repeated here for comparison to Fig S1A.

**Fig S2.**
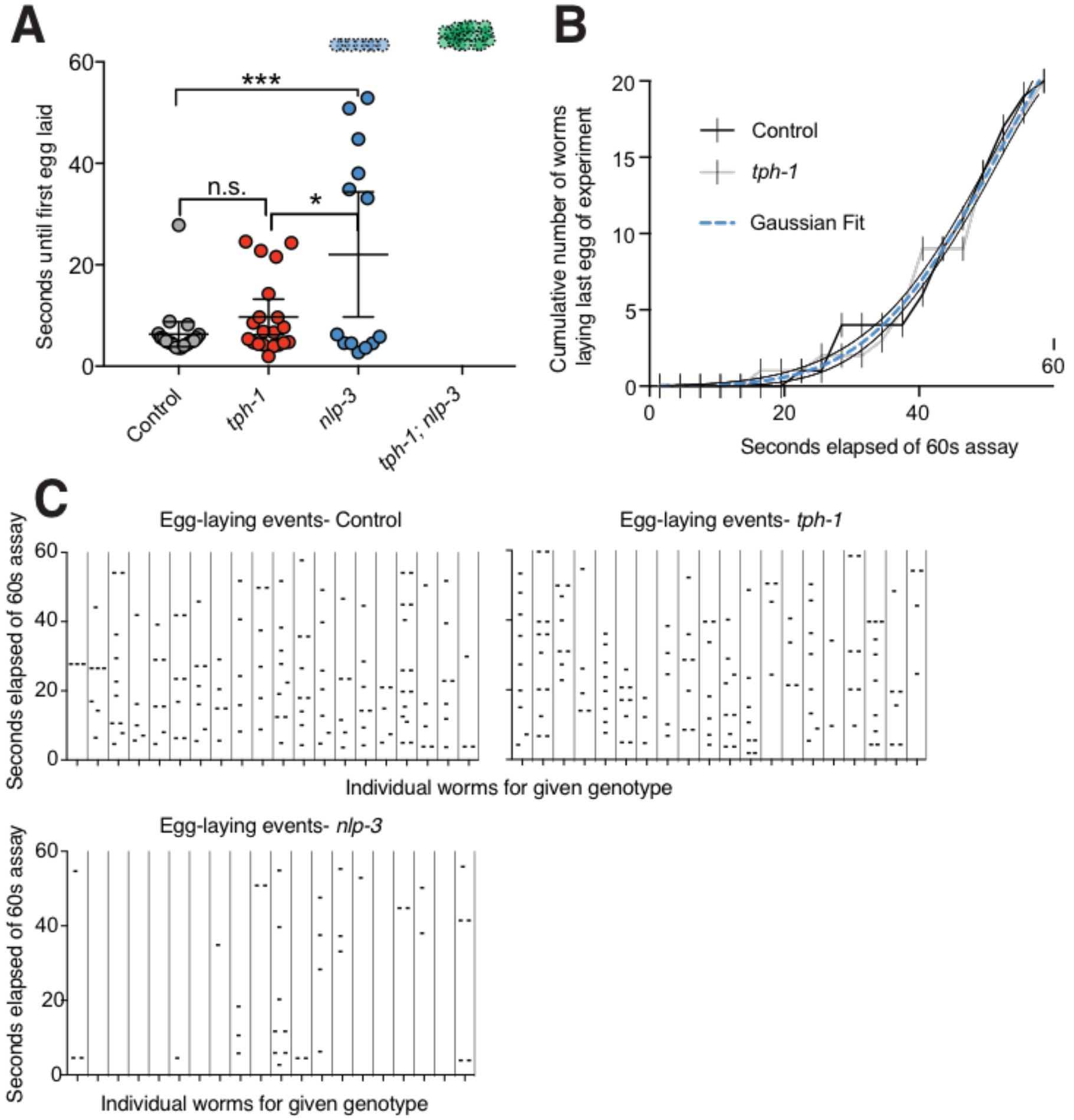
A *tph-1* null mutation does not detectably affect egg laying upon optogenetic activation of HSNs. Measurements of egg-laying upon blue light stimulation of egg-laying in *egl-6p*::ChR2-YFP/+ animals that were controls (wild type for *tph-1* and *nlp-3)* or that carried null mutations in *tph-1* or *nlp-3*. All animals also had a *lite-1(ce314)* mutation that eliminates endogenous responses to blue light. **A)** Average time from onset of blue light simulation to first egg laid, measured as in (73). There was no significant difference between control and *tph-1* animals, but *nlp-3* animals initiated egg laying more slowly and with higher animal-to-animal variability, and 7/20 nlp-3 animals tested failed to lay any eggs. Center line is the mean, error bars are 95% confidence intervals. n.s., no significant difference, *, p<0.033, ***, p<0.001. **B)** Cumulative distribution plot of the time to last egg laid during the 60 second blue light illumination experiment. One Gaussian curve fit both control and *tph-1* data for last egg laid during the assay. **C) Plots of raw dating showing the time point of** each egg laid by each genotype. Each of 20 animals tested per genotype is represented by a vertical column, with each point indicating the time after the onset of blue light illumination when an individual egg was laid Empty columns indicate that no eggs were laid. Two or three horizontally adjacent points indicate eggs laid simultaneously within the 0.05 sec time resolution of our video recording.

**Fig S3.**
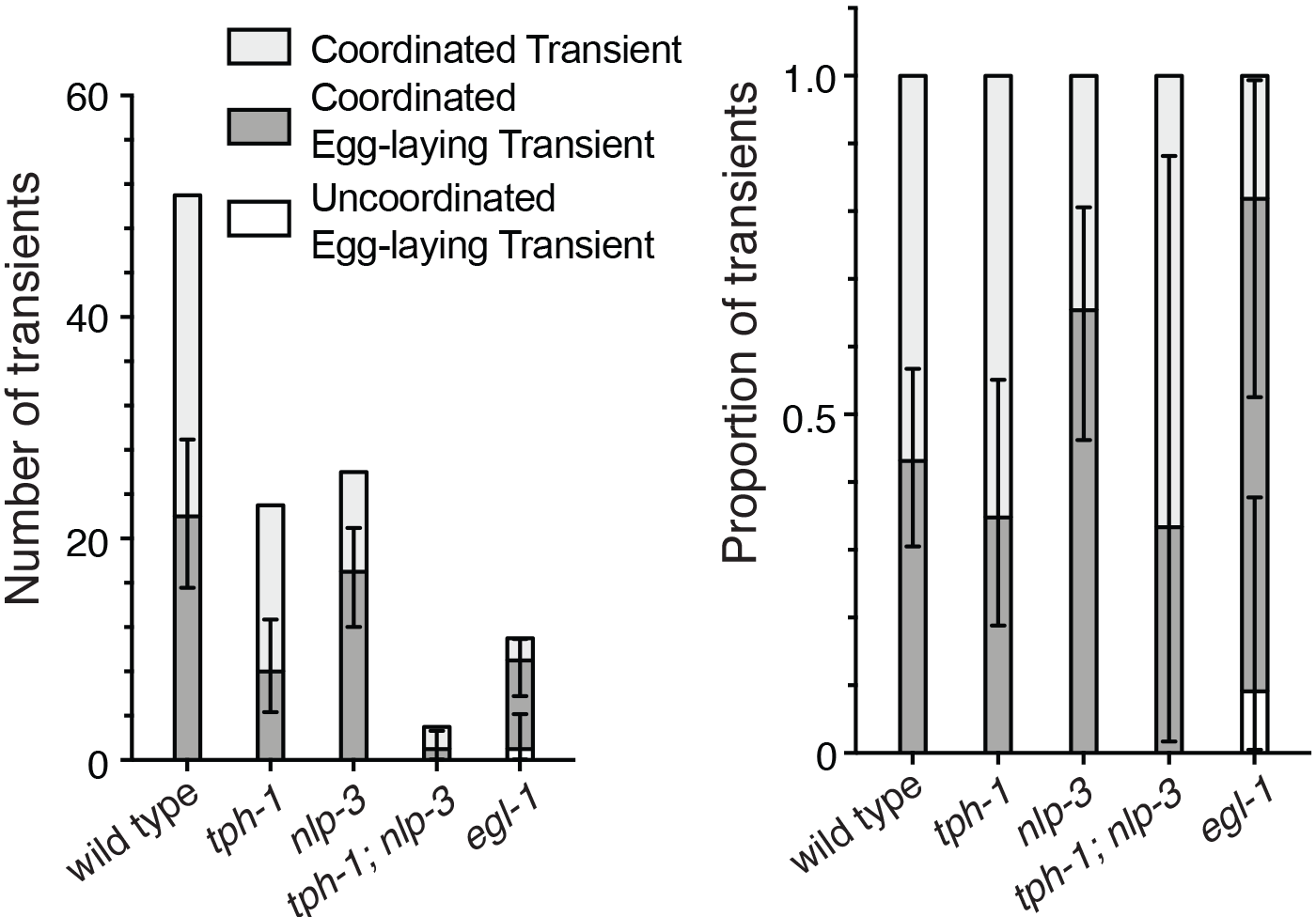
Egg-laying events are associated with coordinated vm1 + vm2 Ca^2+^ transients. The number and proportion of either vm1 or vm1+vm2 coordinated transients that are associated with egg-laying events. One exceptional egg-laying event in an *egl-1* animal occurred after a vm1-only transient, while all 57 others occurred during coordinated vm1+vm2 events. Error bars are 95% confidence intervals for the results expected if an infinite sample size was used.

## References

1. Nautiyal KM, Hen R. Serotonin receptors in depression: from A to B. F1000Res [Internet]. 2017 Feb 9 [cited 2018 Jan 2];6. Available from: https://www.ncbi.nlm.nih.gov/pmc/articles/PMC5302148/

2. Baker KG, Halliday GM, Hornung J-P, Geffen LB, Cotton RGH, To¨rk I. Distribution, morphology and number of monoamine-synthesizing and substance P-containing neurons in the human dorsal raphe nucleus. Neuroscience. 1991;42(3):757–75.

3. Appel NM, Wessendorf MW, Elde RP. Thyrotropin-releasing hormone in spinal cord: coexistence with serotonin and with substance P in fibers and terminals apposing identified preganglionic sympathetic neurons. Brain Research. 1987 Jul 7;415(1):137–43.

4. Henry JN, Manaker S. Colocalization of substance P or enkephalin in serotonergic neuronal afferents to the hypoglossal nucleus in the rat. The Journal of Comparative Neurology. 1998;391(4):491–505.

5. Hökfelt T, Pernow B, Wahren J. Substance P: a pioneer amongst neuropeptides. Journal of Internal Medicine. 2001 Jan 1;249(1):27–40.

6. Nakamura M, Yasuda K, Hasumi-Nakayama Y, Sugiura M, Tomita I, Mori R, et al. Colocalization of serotonin and substance P in the postnatal rat trigeminal motor nucleus and its surroundings. International Journal of Developmental Neuroscience. 2006 Feb;24(1):61–4.

7. Berger M, Gray JA, Roth BL. The Expanded Biology of Serotonin. Annual Review of Medicine. 2009;60(1):355–66.

8. Hornung J-P. The human raphe nuclei and the serotonergic system. Journal of Chemical Neuroanatomy. 2003 Dec 1;26(4):331–43.

9. Sergeyev V, Hökfelt T, Hurd Y. Serotonin and substance P co-exist in dorsal raphe neurons of the human brain. Neuroreport. 1999 Dec 16;10(18):3967–70.

10. Ptak K, Yamanishi T, Aungst J, Milescu LS, Zhang R, Richerson GB, et al. Raphé Neurons Stimulate Respiratory Circuit Activity by Multiple Mechanisms via Endogenously Released Serotonin and Substance P. J Neurosci. 2009 Mar 25;29(12):3720–37.

11. Nusbaum MP, Blitz DM, Marder E. Functional consequences of neuropeptide and small-molecule co-transmission. Nat Rev Neurosci. 2017 Jul;18(7):389–403.

12. Burnstock G. Cotransmission. Current Opinion in Pharmacology. 2004 Feb;4(1):47–52.

13. Burnstock G. Do some nerve cells release more than one transmitter? Neuroscience. 1976 Aug;1(4):239–48.

14. Kupfermann I. Functional studies of cotransmission. Physiological Reviews. 1991 Jul 1;71(3):683–732.

15. Kramer MS, Winokur A, Kelsey J, Preskorn SH, Rothschild AJ, Snavely D, et al. Demonstration of the efficacy and safety of a novel substance P (NK1) receptor antagonist in major depression. Neuropsychopharmacology. 2004 Feb;29(2):385–92.

16. Keller M, Montgomery S, Ball W, Morrison M, Snavely D, Liu G, et al. Lack of Efficacy of the Substance P (Neurokinin1 Receptor) Antagonist Aprepitant in the Treatment of Major Depressive Disorder. Biological Psychiatry. 2006 Feb 1;59(3):216–23.

17. Ratti E, Bellew K, Bettica P, Bryson H, Zamuner S, Archer G, et al. Results From 2 Randomized, Double-Blind, Placebo-Controlled Studies of the Novel NK1 Receptor Antagonist Casopitant in Patients With Major Depressive Disorder. Journal of Clinical Psychopharmacology. 2011 Dec;31(6):727–733.

18. Chalasani SH, Kato S, Albrecht DR, Nakagawa T, Abbott LF, Bargmann CI. Neuropeptide feedback modifies odor-evoked dynamics in Caenorhabditis elegans olfactory neurons. Nat Neurosci. 2010 May;13(5):615–21.

19. Harris G, Mills H, Wragg R, Hapiak V, Castelletto M, Korchnak A, et al. The Monoaminergic Modulation of Sensory-Mediated Aversive Responses in Caenorhabditis elegans Requires Glutamatergic/Peptidergic Cotransmission. J Neurosci. 2010 Jun 9;30(23):7889–99.

20. Schafer WF. Genetics of egg-laying in worms. Annu Rev Genet. 2006;40:487–509.

21. Kupfermann I, Weiss KR. The command neuron concept. Behavioral and Brain Sciences. 1978 Mar;1(1):3–10.

22. Trent C, Tsung N, Horvitz HR. Egg-Laying Defective Mutants of the Nematode Caenorhabditis Elegans. Genetics. 1983 Aug 1;104(4):619–47.

23. Leifer AM, Fang-Yen C, Gershow M, Alkema MJ, Samuel ADT. Optogenetic manipulation of neural activity in freely moving Caenorhabditis elegans. Nat Meth. 2011 Feb;8(2):147–52.

24. Emtage L, Aziz-Zaman S, Padovan-Merhar O, Horvitz HR, Fang-Yen C, Ringstad N. IRK-1 Potassium Channels Mediate Peptidergic Inhibition of Caenorhabditis elegans Serotonin Neurons via a Go Signaling Pathway. J Neurosci. 2012 Nov 14;32(46):16285–95.

25. Collins KM, Bode A, Fernandez RW, Tanis JE, Brewer JC, Creamer MS, et al. Activity of the C. elegans egg-laying behavior circuit is controlled by competing activation and feedback inhibition. eLife Sciences. 2016 Nov 16;5:e21126.

26. Ellis HM, Horvitz HR. Genetic control of programmed cell death in the nematode C. elegans. Cell. 1986 Mar 28;44(6):817–29.

27. Waggoner LE, Zhou GT, Schafer RW, Schafer WR. Control of alternative behavioral states by serotonin in Caenorhabditis elegans. Neuron. 1998 Jul;21(1):203–14.

28. Sze JY, Victor M, Loer C, Shi Y, Ruvkun G. Food and metabolic signalling defects in a Caenorhabditis elegans serotonin-synthesis mutant. Nature. 2000 Feb 3;403(6769):560–4.

29. Zhang Y, Lu H, Bargmann CI. Pathogenic bacteria induce aversive olfactory learning in Caenorhabditis elegans. Nature. 2005 Nov 10;438(7065):179–84.

30. Schafer WF. Genetics of egg-laying in worms. Annu Rev Genet. 2006;40:487–509.

31. Weinshenker D, Garriga G, Thomas JH. Genetic and pharmacological analysis of neurotransmitters controlling egg laying in C. elegans. J Neurosci. 1995 Oct 1;15(10):6975–85.

32. Hobson RJ, Hapiak VM, Xiao H, Buehrer KL, Komuniecki PR, Komuniecki RW. SER-7, a Caenorhabditis elegans 5-HT7-like Receptor, Is Essential for the 5-HT Stimulation of Pharyngeal Pumping and Egg Laying. Genetics. 2006 Jan 1;172(1):159–69.

33. Bany IA, Dong M-Q, Koelle MR. Genetic and Cellular Basis for Acetylcholine Inhibition of Caenorhabditis elegans Egg-Laying Behavior. J Neurosci. 2003 Sep 3;23(22):8060–9.

34. Nathoo AN, Moeller RA, Westlund BA, Hart AC. Identification of neuropeptidelike protein gene families in Caenorhabditis elegans and other species. PNAS. 2001 Nov 20;98(24):14000–5.

35. Kim K, Li C. Expression and regulation of an FMRFamide-related neuropeptide gene family in Caenorhabditis elegans. J Comp Neurol. 2004;475(4):540–50.

36. Altun ZF, Herndon LA, Wolkow CA, Crocker C, Lints R, Hall DH, editors. WormAtlas [Internet]. 2017. Available from: http://www.wormatlas.org

37. Ringstad N, Horvitz HR. FMRFamide neuropeptides and acetylcholine synergistically inhibit egg-laying by C. elegans. Nat Neurosci. 2008 Oct;11(10):1168–76.

38. Chase DL, Koelle MR. Genetic Analysis of RGS Protein Function in Caenorhabditis elegans. In: Methods in Enzymology [Internet]. Academic Press; 2004 [cited 2017 Oct 15]. p. 305–20. (Regulators of G-Protein Signaling, Part A; vol. 389). Available from: http://www.sciencedirect.com/science/article/pii/S0076687904890189

39. Horvitz HR, Chalfie M, Trent C, Sulston JE, Evans PD. Serotonin and octopamine in the nematode Caenorhabditis elegans. Science. 1982 May 28;216(4549):1012–4.

40. Desai C, Garriga G, Mclntire SL, Horvitz HR. A genetic pathway for the development of the Caenorhabditis elegans HSN motor neurons., Published online: 15 December 1988; | doi:101038/336638a0. 1988 Dec 15;336(6200):638–46.

41. Harris G, Mills H, Wragg R, Hapiak V, Castelletto M, Korchnak A, et al. The Monoaminergic Modulation of Sensory-Mediated Aversive Responses in Caenorhabditis elegans Requires Glutamatergic/Peptidergic Cotransmission. J Neurosci. 2010 Jun 9;30(23):7889–99.

42. Carnell L, Illi J, Hong SW, McIntire SL. The G-Protein-Coupled Serotonin Receptor SER-1 Regulates Egg Laying and Male Mating Behaviors in Caenorhabditis elegans. J Neurosci. 2005 Nov 16;25(46):10671–81.

43. Dernovici S, Starc T, Dent JA, Ribeiro P. The serotonin receptor SER-1 (5HT2ce) contributes to the regulation of locomotion in Caenorhabditis elegans. Developmental Neurobiology. 2007;67(2):189–204.

44. Dempsey CM, Mackenzie SM, Gargus A, Blanco G, Sze JY. Serotonin (5HT), fluoxetine, imipramine and dopamine target distinct 5HT receptor signaling to modulate Caenorhabditis elegans egg-laying behavior. Genetics. 2005 Mar;169(3):1425–36.

45. White JG, Southgate E, Thomson JN, Brenner S. The Structure of the Nervous System of the Nematode Caenorhabditis elegans. Phil Trans R Soc Lond B. 1986 Nov 12;314(1165):1–340.

46. Collins KM, Koelle MR. Postsynaptic ERG Potassium Channels Limit Muscle Excitability to Allow Distinct Egg-Laying Behavior States in Caenorhabditis elegans. J Neurosci. 2013 Jan 9;33(2):761–75.

47. Ravi B, Nassar LM, Kopchock RJ, Dhakal P, Scheetz M, Collins KM. Ratiometric Calcium Imaging of Individual Neurons in Behaving Caenorhabditis Elegans. J Vis Exp. 2018 Feb 7;(132).

48. Li P, Collins KM, Koelle MR, Shen K. LIN-12/Notch signaling instructs postsynaptic muscle arm development by regulating UNC-40/DCC and MADD-2 in Caenorhabditis elegans. eLife [Internet]. 2013 Mar 19 [cited 2014 Nov 25];2. Available from: http://www.ncbi.nlm.nih.gov/pmc/articles/PMC3601818/

49. Pearson KG. Common principles of motor control in vertebrates and invertebrates. Annu Rev Neurosci. 1993;16:265–97.

50. Gao S, Guan SA, Fouad AD, Meng J, Kawano T, Huang Y-C, et al. Excitatory motor neurons are local oscillators for backward locomotion. eLife Sciences. 2018 Jan 23;7:e29915.

51. Hapiak VM, Hobson RJ, Hughes L, Smith K, Harris G, Condon C, et al. Dual Excitatory and Inhibitory Serotonergic Inputs Modulate Egg Laying in Caenorhabditis elegans. Genetics. 2009 Jan 1;181(1):153–63.

52. Xiao H, Hapiak VM, Smith KA, Lin L, Hobson RJ, Plenefisch J, et al. SER-1, a Caenorhabditis elegans 5-HT2-like receptor, and a multi-PDZ domain containing protein (MPZ-1) interact in vulval muscle to facilitate serotonin-stimulated egg-laying. Developmental Biology. 2006 Oct 15;298(2):379–91.

53. Gürel G, Gustafson MA, Pepper JS, Horvitz HR, Koelle MR. Receptors and Other Signaling Proteins Required for Serotonin Control of Locomotion in Caenorhabditis elegans. Genetics [Internet]. 2012 Sep 28 [cited 2012 Oct 11]; Available from: http://www.genetics.org/content/early/2012/09/27/genetics.112.142125

54. Fei H, Grygoruk A, Brooks ES, Chen A, Krantz DE. Trafficking of Vesicular Neurotransmitter Transporters. Traffic. 2008 Sep 1;9(9):1425–36.

55. Torrealba F, Carrasco MA. A review on electron microscopy and neurotransmitter systems. Brain Research Reviews. 2004 Dec 1;47(1):5–17.

56. Merighi A. Costorage and coexistence of neuropeptides in the mammalian CNS. Progress in Neurobiology. 2002 Feb 1;66(3):161–90.

57. Mansvelder HD, Kits KS. Calcium channels and the release of large dense core vesicles from neuroendocrine cells: spatial organization and functional coupling. Progress in Neurobiology. 2000 Nov 1;62(4):427–41.

58. M G, Eg A, O H. A cellular and regulatory map of the GABAergic nervous system of C. elegans., A cellular and regulatory map of the GABAergic nervous system of C. elegans. Elife [Internet]. 2016 [cited 2017 Jul 21];5, 5. Available from: http://europepmc.org/abstract/MED/27740909, http://europepmc.org/articles/PMC5065314/?report=abstract

59. Ritter AD, Shen Y, Fuxman Bass J, Jeyaraj S, Deplancke B, Mukhopadhyay A, et al. Complex expression dynamics and robustness in C. elegans insulin networks. Genome Res. 2013 Jun;23(6):954–65.

60. Croset V, Treiber CD, Waddell S. Cellular diversity in the Drosophila midbrain revealed by single-cell transcriptomics. eLife Sciences. 2018 Apr 19;7:e34550.

61. Nassel DR. Substrates for Neuronal Cotransmission With Neuropeptides and Small Molecule Neurotransmitters in Drosophila. Front Cell Neurosci [Internet]. 2018 [cited 2018 May 14];12. Available from: https://www.frontiersin.org/articles/10.3389/fncel.2018.00083/full

62. Hökfelt T, Millhorn D, Seroogy K, Tsuruo Y, Ceccatelli S, Lindh B, et al. Coexistence of peptides with classical neurotransmitters. Experientia. 1987 Jul 1;43(7):768–80.

63. Marder E. Neuromodulation of Neuronal Circuits: Back to the Future. Neuron. 2012 Oct 4;76(1):1–11.

64. Marder E. Neuromodulation of Neuronal Circuits: Back to the Future. Neuron. 2012 Oct 4;76(1):1–11.

65. Leinwand SG, Chalasani SH. Neuropeptide signaling remodels chemosensory circuit composition in Caenorhabditis elegans. Nat Neurosci. 2013 Oct;16(10):1461–7.

66. Brenner S. The Genetics of Caenorhabditis Elegans. Genetics. 1974 May 1;77(1):71–94.

67. Evans T. Transformation and microinjection. WormBook [Internet]. 2006 [cited 2017 Oct 15]; Available from: http://www.wormbook.org/chapters/www_transformationmicroinjection/transformationmicroinjection.html

68. Fay DS. Classical genetic methods. WormBook. 2013 Dec 30;1–58.

69. Bhatla N, Droste R, Sando SR, Huang A, Horvitz HR. Distinct Neural Circuits Control Rhythm Inhibition and Spitting by the Myogenic Pharynx of C. elegans. Current Biology. 2015 Aug 17;25(16):2075–89.

70. Sze JY, Victor M, Loer C, Shi Y, Ruvkun G. Food and metabolic signalling defects in a Caenorhabditis elegans serotonin-synthesis mutant. Nature. 2000 Feb 3;403(6769):560–4.

71. Chase DL, Koelle MR. Genetic Analysis of RGS Protein Function in Caenorhabditis elegans. In: Methods in Enzymology [Internet]. Academic Press; 2004 [cited 2017 Oct 15]. p. 305–20. Available from: http://www.sciencedirect.com/science/article/pii/S0076687904890189

72. Leifer AM, Fang-Yen C, Gershow M, Alkema MJ, Samuel ADT. Optogenetic manipulation of neural activity in freely moving Caenorhabditis elegans. Nat Meth. 2011 Feb;8(2):147–52.

73. Emtage L, Aziz-Zaman S, Padovan-Merhar O, Horvitz HR, Fang-Yen C, Ringstad N. IRK-1 Potassium Channels Mediate Peptidergic Inhibition of Caenorhabditis elegans Serotonin Neurons via a Go Signaling Pathway. J Neurosci. 2012 Nov 14;32(46):16285–95.

74. C. elegans Fosmid Library [Internet]. [cited 2017 Oct 15]. Available from: https://www.sourcebioscience.com/products/life-science-research/clones/genomic-clones/c-elegans-genomic-clone-collections/c-elegans-fosmid-library/

75. Perkins J, Wong K, Warren R, Schein J, Stott J, Holt R, et al. A Caenorhabditis elegans fosmid library. In: International Worm Meeting [Internet]. 2005 [cited 2017 Oct 15]. Available from: http://www.wormbase.org/db/misc/paper?name=WBPaper00026350

76. Chase DL, Koelle MR. Genetic Analysis of RGS Protein Function in Caenorhabditis elegans. In: Methods in Enzymology [Internet]. Academic Press; 2004 [cited 2018 Jan 4]. p. 305–20. (Regulators of G-Protein Signaling, Part A; vol. 389). Available from: http://www.sciencedirect.com/science/article/pii/S0076687904890189

77. Edwards SL, Charlie NK, Milfort MC, Brown BS, Gravlin CN, Knecht JE, et al. A Novel Molecular Solution for Ultraviolet Light Detection in Caenorhabditis elegans. PLOS Biology. 2008 Aug 5;6(8):e198.

78. Edwards SL, Charlie NK, Milfort MC, Brown BS, Gravlin CN, Knecht JE, et al. A Novel Molecular Solution for Ultraviolet Light Detection in Caenorhabditis elegans. PLOS Biology. 2008 Aug 5;6(8):e198.

